# Establishment of spinocerebellar ataxia type 34 model mice accompanied by early glial activation and degeneration of cerebellar neurons

**DOI:** 10.1101/2025.10.15.682296

**Authors:** Yuri Morikawa-Yujiri, Kensuke Motomura, Ayumu Konno, Natsuko Hitora-Imamura, Yuki Kurauchi, Satohiro Masuda, Hirokazu Hirai, Hiroshi Katsuki, Takahiro Seki

## Abstract

Spinocerebellar ataxia type 34 (SCA34) is an autosomal dominant neurodegenerative disease primarily characterized by progressive cerebellar atrophy and ataxia, frequently accompanied by cognitive dysfunction and erythrokeratodermia variabilis. In 2014, missense mutations in the gene encoding elongation of very long chain fatty acids protein 4 (ELOVL4) were identified as the causative gene for SCA34. ELOVL4, which is involved in the synthesis of very long chain fatty acids, is highly expressed in the cerebellum compared to other brain regions, with predominant expression in neurons. We attempted to establish a mouse model of SCA34 by expressing mutant ELOVL4 in cerebellar neurons using adeno-associated virus (AAV) vectors and to elucidate the underlying pathogenic mechanisms. Expression of W246G mutant ELOVL4 successfully induced progressive motor dysfunction beginning at two weeks post-AAV vector injection. Immunohistochemical analyses revealed that the degeneration of cerebellar Purkinje cells and neurons in the deep cerebellar nuclei (DCN) paralleled the observed motor decline. Importantly, microglial activation was detected in the molecular layer of the cerebellar cortices and the DCN prior to the onset of both neurodegeneration and motor dysfunction. Furthermore, after the onset of motor symptoms, the SCA34 model mice exhibited decreased synaptic inputs from climbing fibers to Purkinje cells, as well as reduced inputs from Purkinje cells to DCN neurons. These findings suggest that early microglial activation and the resulting synaptic disturbance are critical preceding events that lead to the progressive cerebellar neurodegeneration and motor dysfunction observed in this SCA34 mouse model.

## 1. Introduction

Spinocerebellar ataxias (SCAs) are a disease group of autosomal-dominant neurodegenerative diseases primarily characterized by progressive cerebellar atrophy and ataxia. Currently, SCAs are classified into type 1 through 51 based on their respective causative genes (Klockgether et al., 2019; Sullivan et al., 2018). While six SCA types are caused by expanded CAG repeats that translate into polyglutamine chains, the other types arise from the missense or nonsense mutation in various causative genes. The focus of the present study is spinocerebellar ataxia type 34 (SCA34). In addition to cerebellar ataxia, SCA34 patients often present with other symptoms, including cognitive impairment and erythrokeratodermia variabilis (Haeri et al., 2021; Kirola et al., 2022; Ozaki et al., 2019). In 2014, missense mutations in the gene encoding elongation of very long chain fatty acids protein 4 (ELOVL4) were identified as the causative factor for SCA34 (Cadieux-Dion et al., 2014). ELOVL4 is an elongase localized to the endoplasmic reticulum membrane (Jakobsson et al., 2006) that is critical for the elongation of very long-chain (VLC) polyunsaturated and saturated fatty acids (C28 and above) (Agbaga et al., 2008). ELOVL4 is highly expressed in the retina, skin, thymus, and brain (Leonard et al., 2004). Within the brain, its expression is predominantly and most abundant in cerebellar neurons (Sherry et al., 2017). To date, six missense mutations (I171T, L168F, L168S, Q180P, T233M, W246G) have been identified in SCA34 patients (Nishide et al., 2024). In vitro studies have shown that these SCA34 mutations impair ELOVL4’s ability to elongate and generate VLC-saturated fatty acids (Gyening et al., 2023).

Postmortem neuropathological examinations of SCA34 patients have consistently revealed a loss of cerebellar Purkinje cells and shrinkage of the molecular layer (ML) in cerebellar cortices (Ellezam et al., 2023; Ozaki et al., 2021a). However, a recently developed knock-in rat model carrying the W246G mutant ELOVL4 exhibited progressive motor dysfunction but lacked the characteristic degeneration of Purkinje cells or other cerebellar neurons (Fessler et al., 2024). This rat model did show dysregulated synaptic transmission and plasticity (Nagaraja et al., 2023, 2021). To address the need for a mouse model that recapitulates the major pathological hallmarks observed in SCA34 patients, we constructed SCA34 model mice via adeno-associated viral (AAV) vector-mediated gene transfer of the W246G mutant ELOVL4 into cerebellar neurons. These mice developed early-onset progressive ataxia accompanied by Purkinje cell degeneration and ML shrinkage, thus sharing the key pathological features with SCA34 patients. Furthermore, these mice exhibited shrinkage and reduction in neurons of the deep cerebellar nuclei (DCN), the primary targets of Purkinje cell axons. Our novel SCA34 model mice will serve as a valuable tool for elucidating the precise mechanisms by which mutant ELOVL4 triggers cerebellar dysfunction and for screening potential therapeutic or preventive agents against SCA34.

## 2. Materials and methods

### 2.1. Materials

Anti-DYKDDDDK (FLAG) mouse monoclonal antibody (clone 2H8, catalog No. KO602) was obtained from TransGenic Group (Fukuoka, Japan). Rabbit polyclonal antibodies against protein kinase Cγ (PKCγ, Catalog No. GTX107639) and vesicular GABA transporter (VGAT, Catalog No. GTX101908) were obtained from GeneTex (Irvine, CA, USA). Anti-ionized calcium-binding adapter molecule 1 (Iba1) rabbit polyclonal antibody (Catalog No. 019−19741) was obtained from FUJIFILM Wako Pure Chemical Industries (Osaka, Japan). Guinea pic polyclonal antibodies against vesicular glutamate transporter 1 (VGluT1, Catalog No. AB5905) and vesicular glutamate transporter 2 (VGluT2, Catalog No. AB2551) were obtained from Merck Millipore (Burlington, MA, USA). Rabbit monoclonal antibodies against glial fibrillary acidic protein (GFAP, Catalog No. 12389), calbindin D28k (Catalog No. 1317), and-MAP2 (Catalog No. 8707) antibodies were obtained from Cell Signaling Technologies (Danvers, MA, USA). The following donkey polyclonal secondary antibodies conjugated to Alexa Fluor were purchased from Thermo Fisher Scientific: Alexa Fluor 488-conjugated anti-mouse IgG (Cat. No. A−21202), Alexa Fluor 555-conjugated anti-mouse IgG (Cat. No. A-31570), Alexa Fluor 555-conjugated anti-rabbit IgG (Cat. No. A−31572), and Alexa Fluor 488-conjugated anti-guinea pig IgG (Cat. No. A−11073).

### 2.2. Ethical approval

All animal experiments involving mice were performed in accordance with the Guidelines of the United States National Institutes of Health for the care and use of animals for experimental procedures and were approved by the Kumamoto University Ethics Committee concerning animal experiments (Approval No. A2025-007). Experiments concerning AAV vectors were conducted under the Cartagena Protocol on Biosafety to the Convention on Biological Diversity and were approved by the Kumamoto University Ethics Committee concerning experiments on genetically modified organisms (Approval No. 5-016).

### 2.3. AAV vector injection to the mouse cerebellum

cDNAs encoding N-terminally FLAG-tagged wild-type (WT) and SCA34 mutant (W246G) ELOVL4 (FLAG-ELOVL4) were generated using a PrimeStar mutagenesis basal kit (Catalog No. R046A, Takara Bio, Kusatsu, Japan). These cDNAs were subcloned into the expression vector for the construction of AAV vectors. We constructed AAV serotype 9 (AAV9) vectors to express WT or W246G mutant FLAG-ELOVL4 under the neuron-specific synapsin I promoter as previously described (Huda et al., 2014). We also constructed an AAV9 vector to express green fluorescent protein (GFP) under the Purkinje cell-specific L7 promoter for visualizing Purkinje cell axons.

For the construction of SCA34 model mice, 4-week-old male ICR mice were stereotaxically injected with AAV9 vectors (3.2 × 10^9^ vector genomes, vg) to express WT or W246G mutant FLAG-ELOVL4 into the cerebellum (Ohta et al., 2021). Mice were anesthetized with a mixture of 3 mg/kg medetomidine hydrochloride, 4 mg/kg midazolam, and 5 mg/kg butorphanol tartrate. The AAV vector solution (8 μL at a concentration of 4×10^11^ vg/mL) was injected into the cerebellar parenchyma over 30 min at a constant rate using a microinfusion pump, guided by stereotaxic coordinates: 6.5 mm caudal to the bregma and 1.9 mm below the skull. Control mice received an equal volume (8 μL) of phosphate-buffered saline (PBS). In several experiments, an equal amount of AAV9 vectors to express GFP was co-injected with vectors to express FLAG-ELOVL4.

### 2.4. Assessment of motor function

Motor function was assessed by the beam-walking test (Hijioka et al., 2017) at 1, 2, 4, 6, and 8 weeks following AAV vector injection. The motor performance of mice was quantified by measuring two parameters: fault rates, which represented the frequency of hind-limb slips while traversing the beam (1.1 m length, 1.5 cm width, and 50 cm height), and walking speed.

### 2.5. Histological analyses of cerebellar sections

Mice were sacrificed and fixed by transcardial perfusion with 4% paraformaldehyde (PFA) at 1, 2, and 8 weeks post-AAV vector injection. After post-fixation, 30 μm-thick coronal cerebellar sections were prepared for immunofluorescence analysis. Immunofluorescence was performed as described previously (Hijioka et al., 2017) using following primary antibodies: anti-DYKDDDDK (FLAG) (1:1,000), anti-PKCγ (1:500), anti-MAP2 (1:500), anti-GFAP (1:500), anti-Iba1 (1:500), anti-VGluT1 (1:500), anti-VGlut2 (1:500), anti-VGAT (1:500), anti-calbindin (1:500), and anti-NeuN (1:500). Following incubation with Alexa Fluor 488-or Alexa Fluor 555-conjuugated secondary antibodies, fluorescence images of cerebellar slices were captured using fluorescence microscopes: BZ-9000 (Keyence, Osaka, Japan) and THUNDER imager (Leica Biosystems, Nussloch, Germany).

For the quantification of protein expression, immunofluorescence intensities were measured from the acquired images using ImageJ software (National Institute of Health, Bethesda, MD, USA). The relative fluorescence intensity was quantified by normalizing the intensity within the expression area to the intensity of areas where FLAG-ELOVL4 was not expressed. The averaged intensity of 2-3 images per mouse was used as the quantified datum for each mouse.

### 2.6. Statistical analyses

All quantitative data are represented as the mean ± standard error of the mean (SEM). The results of the beam-walking test were analyzed using two-way ANOVA followed by a Bonferroni post hoc test. The results of immunofluorescence experiments were analyzed using one-way ANOVA followed by a Tukey’s post hoc test. Statistical analyses were performed using the GraphPad Prism six software (GraphPad Software, San Diego, CA, United States). Probability values (p) less than 0.05 were considered statistically significant.

## 3. Results

### 3.1. Expression of SCA34 mutant ELOVL4 in cerebellar neurons induces progressive motor dysfunction

In the cerebellum, endogenous ELOVL4 is primarily restricted to neurons including Purkinje cells, granule cells, and interneurons in the ML (Sherry et al., 2017). Therefore, we expressed N-terminally FLAG-tagged wild-type (WT) or SCA34 mutant (W246G) ELOVL4 (FLAG-ELOVL4) in cerebellar neurons using AAV9 vectors driven by a neuron-specific synapsin I promoter. As previously reported (Ohta et al., 2021), we injected either PBS or AAV vectors into the cerebellar white matter. The motor function of mice was evaluated by the beam walking test at 1, 2, 4, 6 and 8 weeks following the cerebellar injection (Fig. 1A). Compared with PBS-injected mice, mice expressing WT FLAG-ELOVL4 showed mild motor impairment, characterized by a trend forwards an elevated fault rate and a significant reduction in walking speed starting at 4 weeks post-injection (Fig. 1B, C). In sharp contrast, mice expressing the W246G mutant FLAG-ELOVL4 exhibited an earlier onset and more severe motor dysfunction, characterized by a progressive and significant increase in fault rate and a decline in walking speed starting at 2 weeks post-injection (Fig. 1B, C). These findings indicate that mice expressing mutant ELOVL4 in the cerebellar neurons could serve as model mice for SCA34, accompanied by early progressive motor dysfunction.

**Figure 1.**
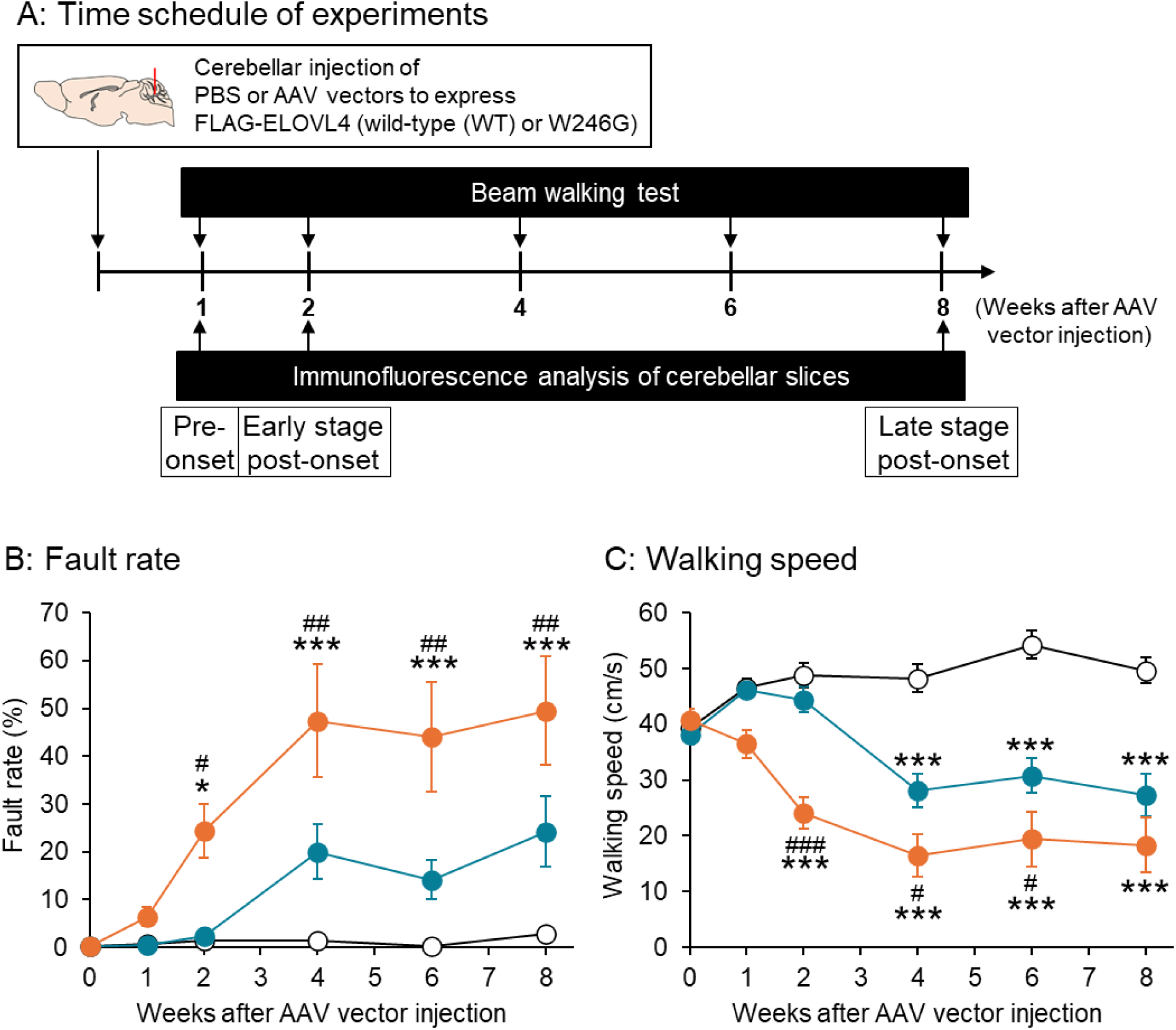
Motor performance of mice expressing wild-type and SCA34 mutant FLAG-ELOVL4 in cerebellar neurons. A. Time schedule of experiments. B, C. Temporal changes in fault rates (B) and walking speed (C) during beam walking tests. Data are shown for PBS-injected control mice (white), WT FLAG-ELOVL4-expressing mice (WT mice, blue), and W246G mutant FLAG-ELOVL4-expressing mice (W246G mice, orange). * p < 0.05, *** p < 0.001 vs control mice; # p < 0.05, ## p < 0.01, ### p < 0.001 vs WT mice (Two-way ANOVA, n = 11 for controls, n = 12 for WT, and n = 15 for W246G).

### 3.2. SCA34 mutant ELOVL4 causes progressive neurodegeneration of cerebellar Purkinje cells and neurons in the deep cerebellar nuclei

Next, we investigated the effects of the expression of WT and W246G mutant FLAG-ELOVL4 on the morphology and survival of neurons in the cerebellar cortices using immunofluorescence experiments at three different time points: 1 week (pre-onset of motor dysfunction), 2 weeks (early-stage post-onset), and 8 weeks (late-stage post-onset) after the AAV vector injection (Fig. 1A). Immunofluorescence studies with anti-PKCγ and MAP2 antibodies, which label Purkinje cells and cerebellar neurons (including Purkinje cells, and granule cells, and ML interneurons), respectively, revealed that FLAG-ELOVL4 was primarily expressed in Purkinje cells and ML interneurons, and partially in neurons of the granule cell layer in the cerebellar cortices (Fig. 2). Pre-onset (1 week): WT and W246G mutant FLAG-ELOVL4 did not affect the Purkinje cell count or ML thickness (Fig. 2A-D). Early-stage post-onset (2 weeks): While WT ELOVL4 did not induce morphological changes or impact of the survival of cerebellar neurons, the W246G mutant ELOVL4 triggered a prominent loss of Purkinje cells and a subtle decrease in ML thickness (Fig. 2E-H). Late-stage post-onset (8 weeks): when both WT and W246G mutant FLAG-ELOVL4 groups displayed a significant decline in motor function, both groups showed a significant decrease in the Purkinje cell count and ML thickness (Fig. 2I-L).

**Figure 2.**
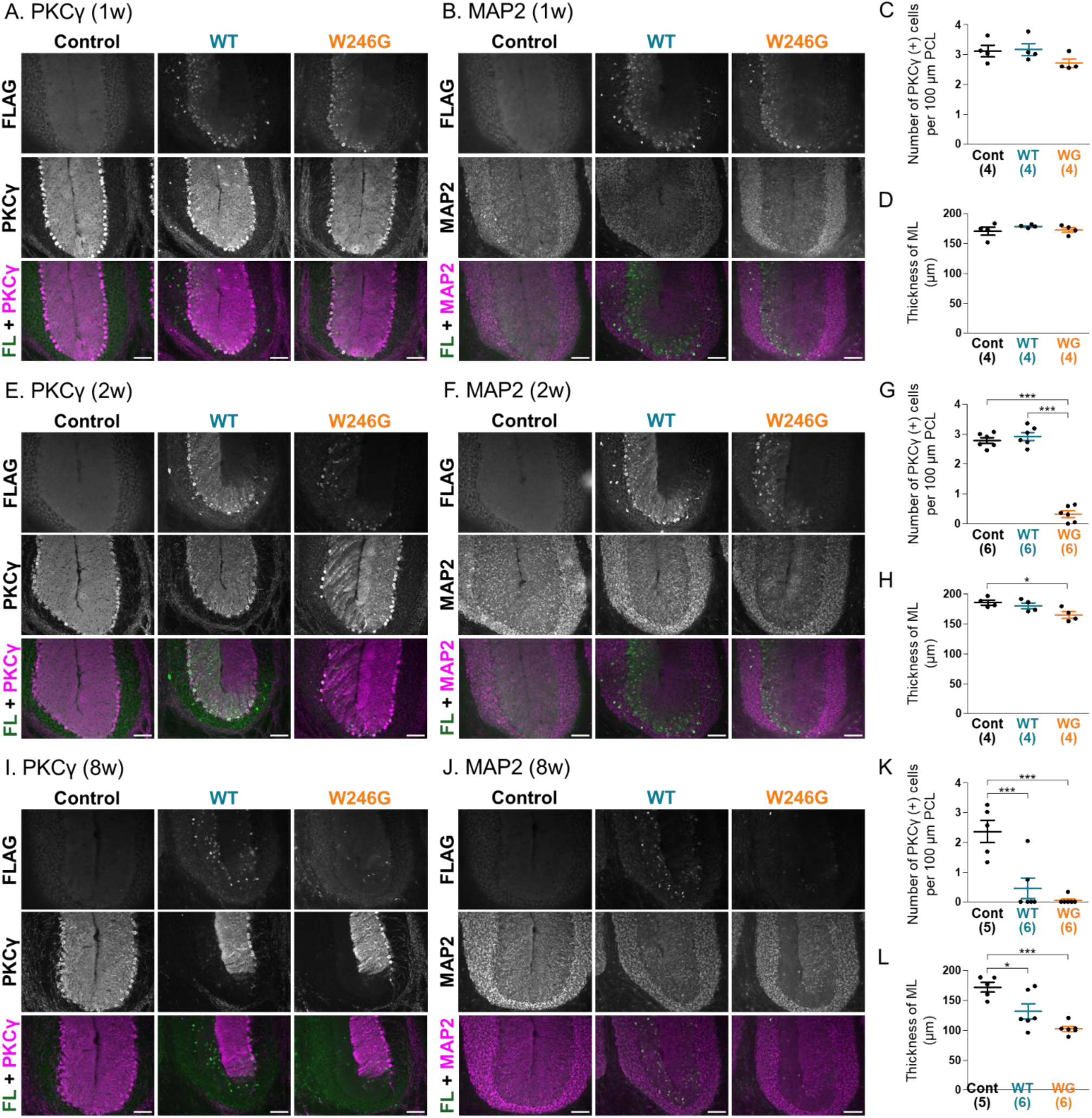
Histological changes in neurons of the cerebellar cortex in mice expressing wild-type and SCA34 mutant FLAG-ELOVL4 in cerebellar neurons. A,E,I. Representative immunofluorescence images showing FLAG, PKCγ, and merged FLAG + PKCγ expression in the cerebellar cortex of control, WT, and W246G mice at 1 week (pre-onset of motor dysfunction, A), 2 weeks (early-stage post-onset, E), and 8 weeks (late-stage post-onset, I) after AAV vector injection. B,F,J. Representative immunofluorescence images showing FLAG, MAP2, and merged FLAG + MAP2 expression in the cerebellar cortex of control, WT, and W246G mice at 1 week (B), 2 weeks (F), and 8 weeks (J) after AAV vector injection. Scale bars are 100 μm. C,G,K. Quantitative analysis of the number of PKCγ-positive Purkinje cells at 1 week (C), 2 weeks (G), and 8 weeks (K) post-injection. The cell count is presented as the number of Purkinje cell somata per 100 μm length of the Purkinje cell layer. D,H,L. Quantitative analysis of the thickness of the molecular layer (ML) at 1 week (D), 2 weeks (H), and 8 weeks (L) post-injection. Numbers in parentheses in the graphs represent the number of mice assessed per group. * p < 0.05, *** p < 0.001 (One-way ANOVA).

We also examine the effects of WT and W246G mutant ELOVL4 expression in the deep cerebellar nuclei (DCN), which receive inhibitory synaptic inputs from Purkinje cells (Watson et al., 2015). Both WT and W246G mutant FLAG-ELOVL4 were expressed in the MAP2-positive DCN neurons (Fig. 3). Pre-onset (1 week): WT and W246G mutant ELOVL4 did not affect the somatic size or the number of MAP2-positive DCN neurons (Fig. 3A-C). Early-stage post-onset (2 weeks): The W246G mutant ELOLV4 significantly reduced the somatic size and the number of DCN neurons, while WT ELOVL4 had no effect (Fig. 3D-F). Late-stage post-onset (8 weeks): WT ELOVL4 also induced a reduction in the somatic size of DCN neurons. However, the W246G mutant ELOVL4 caused even greater decreases in both the somatic size and the number of DCN neurons (Fig. 3G-I). These findings indicate that overexpression of both WT and W246G mutant FLAG-ELOVL4 causes progressive damage to Purkinje cells and DCN neurons, which occurs in parallel with the onset and progression of motor dysfunction.

**Figure 3.**
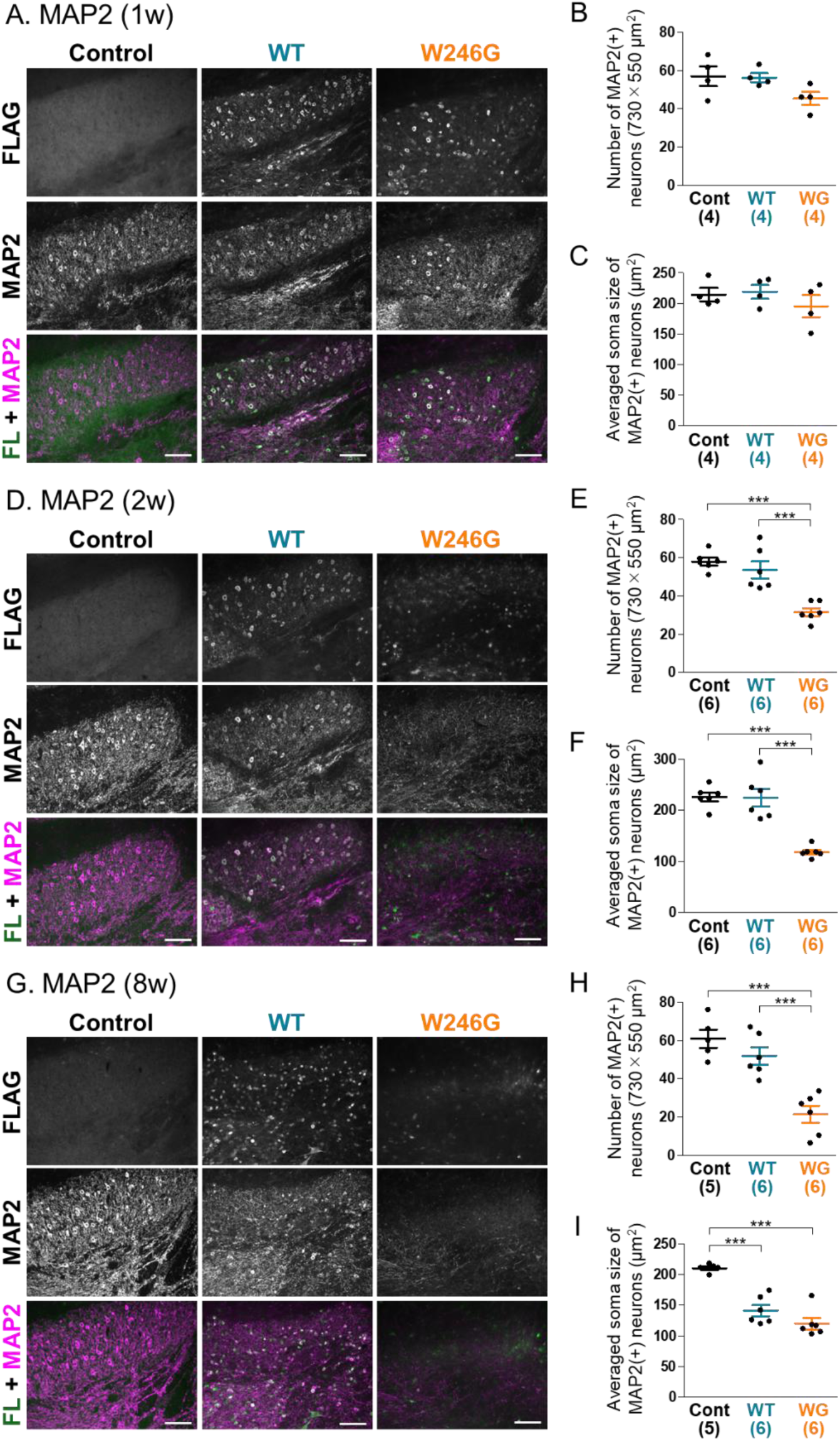
Histological changes in neurons of the deep cerebellar nuclei (DCN) in mice expressing wild-type and SCA34 mutant FLAG-ELOVL4 in cerebellar neurons. A,D,G. Representative immunofluorescence images showing FLAG, MAP2, and merged FLAG + MAP2 expression in the DCN of control, WT, and W246G mice at 1 week (A), 2 weeks (D), and 8 weeks (F) after AAV vector injection. Scale bars are 100 μm. B,E,H. Quantitative analysis of the number of MAP2-positive DCN neurons at 1 week (B), 2 weeks (E), and 8 weeks (H) post-injection. DCN neurons were counted in the same areas shown in A, D, and G. C,F,I. Quantitative analysis of the average soma size of MAP2-positive DCN neurons at 1 week (C), 2 weeks (F), and 8 weeks (I) post-injection. Numbers in parentheses in the graphs represent the number of mice assessed per group. *** p < 0.001 (One-way ANOVA).

### 3.3. Expression of SCA34 mutant ELOVL4 in cerebellar neurons triggers early activation of microglia, followed by the activation of astrocytes

We next investigated the temporal changes in the activation of astrocytes and microglia in the ML of the cerebellar cortices and DCN. We used glial fibrillary acidic protein (GFAP) and ionized calcium-binding adapter molecule 1 (Iba1) as markers for astrocytes and microglia, respectively. In the ML, GFAP and Iba1 expression levels were significantly elevated in mice expressing WT FLAG-ELOVL4 at the late-stage post-onset (Fig. 4I-L). In contrast, in mice expressing W246G mutant ELOVL4, the GFAP level in the ML significantly increased at both the early-and late-stages post-onset, while the Iba1 level in the ML significantly increased at all time points examined, including the presymptomatic stage (Fig. 4).

**Figure 4.**
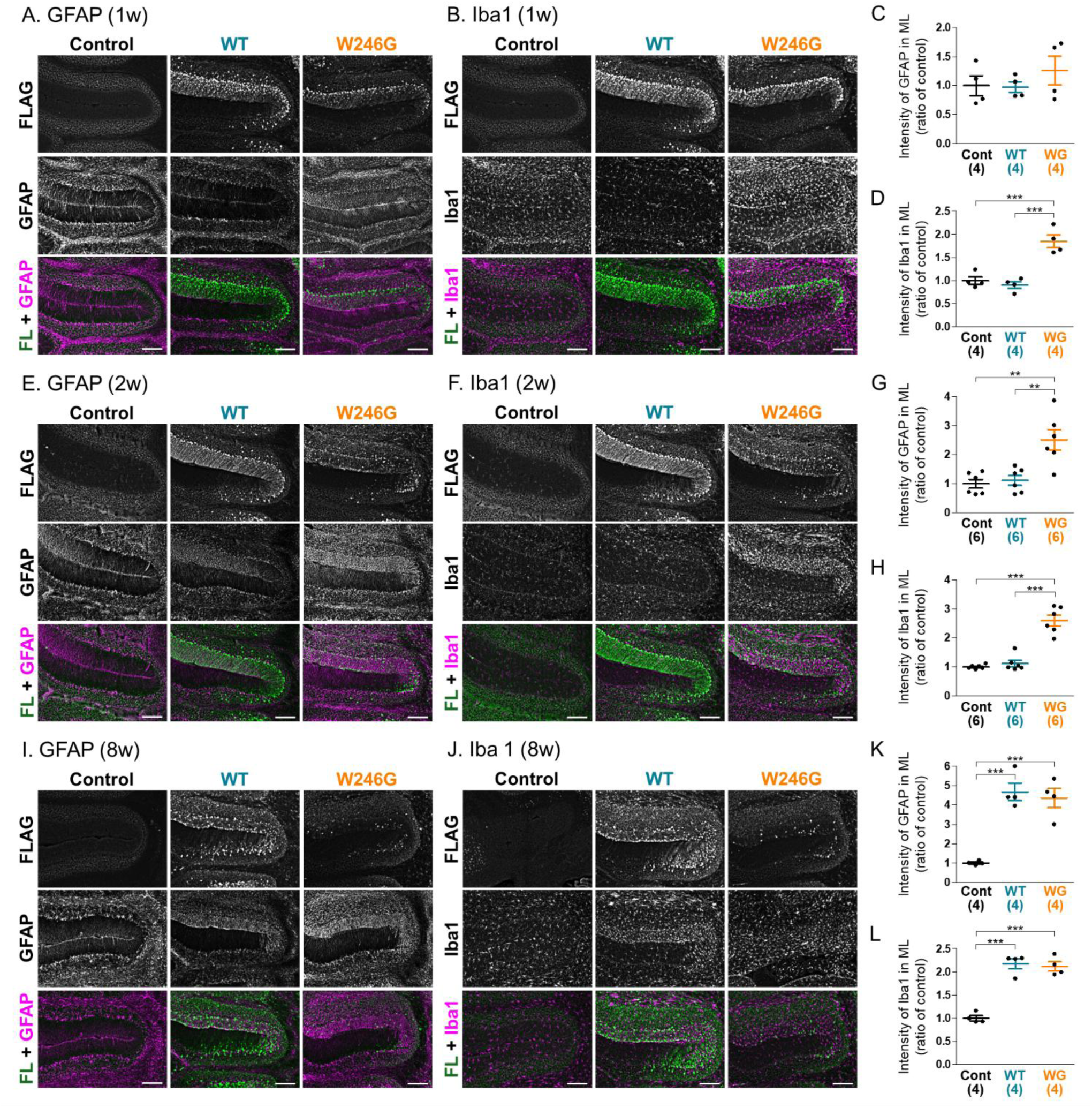
Histological changes in glial cells of the cerebellar cortex in mice expressing wild-type and SCA34 mutant FLAG-ELOVL4 in cerebellar neurons. A,E,I. Representative immunofluorescence images showing FLAG, GFAP, and merged FLAG + GFAP expression in the cerebellar cortex of control, WT, and W246G mice at 1 week (A), 2 weeks (E), and 8 weeks (I) after AAV vector injection. B,F,J. Representative immunofluorescence images showing FLAG, Iba1, and merged FLAG + Iba1 expression in the cerebellar cortex of control, WT, and W246G mice at 1 week (B), 2 weeks (F), and 8 weeks (J) after AAV vector injection. Scale bars are 100 μm. C,G,K. Quantitative analysis of GFAP intensity in the ML at 1 week (C), 2 weeks (G), and 8 weeks (K) post-injection. D,H,L. Quantitative analysis of Iba1 intensity in the ML at 1 week (D), 2 weeks (H), and 8 weeks (L) post-injection. Numbers in parentheses in the graphs represent the number of mice assessed per group. ** p < 0.01, *** p < 0.001 (One-way ANOVA).

Regarding the DCN, the GFAP level was not significantly affected at any time point in mice expressing WT ELOVL4 (Fig. 5). The Iba1 level in the DCN was significantly elevated in mice expressing WT ELOVL4 at the late-stage post-onset (Fig. 5J,L). In mice expressing W246G mutant ELOVL4, the alterations of GFAP and Iba1 levels in DCN (Fig. 5) mirrored those observed in the ML (Fig.4). These findings revealed that microglial activation of microglia in both the ML and the DCN occurred prior to the onset of the motor dysfunction in the SCA34 model mice.

**Figure 5.**
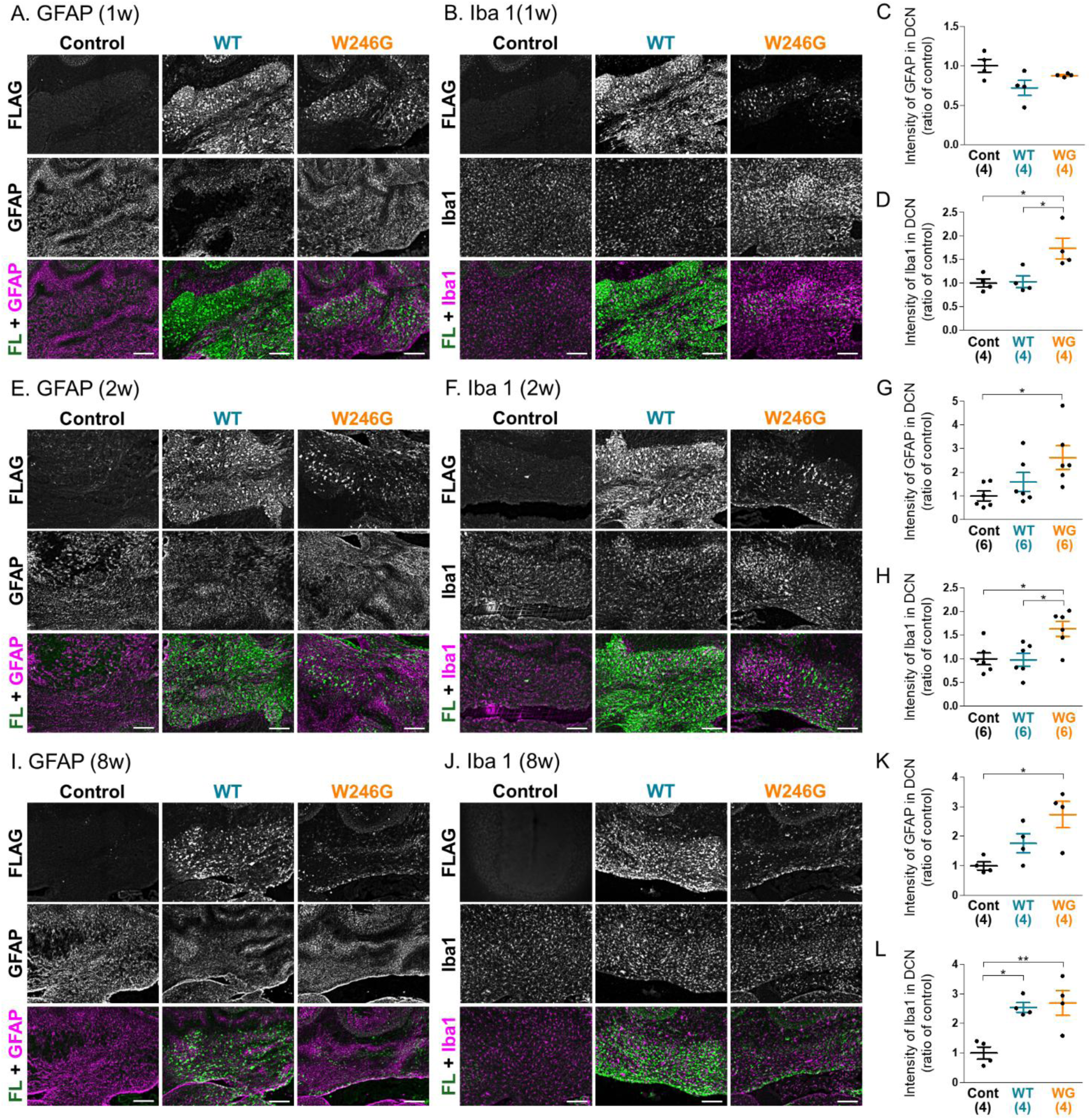
Histological changes in glial cells of the DCN in mice expressing wild-type and SCA34 mutant FLAG-ELOVL4 in cerebellar neurons. A,E,I. Representative immunofluorescence images showing FLAG, GFAP, and merged FLAG + GFAP expression in the DCN of control, WT, and W246G mice at 1 week (A), 2 weeks (E), and 8 weeks (I) after AAV vector injection. B,F,J. Representative immunofluorescence images showing Iba1, FLAG, and merged Iba1 + FLAG expression in the DCN of control, WT, and W246G mice at 1 week (B), 2 weeks (F), and 8 weeks (J) after AAV vector injection. Scale bars are 100 μm. C,G,K. Quantitative analysis of GFAP intensity in the DCN at 1 week (C), 2 weeks (G), and 8 weeks (K) post-injection. D,H,L. Quantitative analysis of Iba1 intensity in the DCN at 1 week (D), 2 weeks (H), and 8 weeks (L) post-injection. Numbers in parentheses in the graphs represent the number of mice assessed per group. * p < 0.05, ** p < 0.01 (One-way ANOVA).

### 3.4. SCA34 mutant ELOVL4 reduces synaptic inputs from climbing fibers to Purkinje cells and from Purkinje cells to neurons in the deep cerebellar nuclei

Next, we investigated the effects of FLAG-ELOVL4 expression on excitatory synaptic inputs to Purkinje cells. In the ML, vesicular glutamate transporter 1 (VGluT1) and vesicular glutamate transporter 2 (VGluT2) are localized at the presynaptic terminals of the parallel and climbing fibers, respectively, which terminate on the dendrites of Purkinje cells (Hioki et al., 2003). The expression VGluT1 level in the ML of mice expressing WT FLAG-ELOVL4 was unaffected at the early-and late-stages post-onset. However, the level of VGluT1 in the ML of mice expressing W246G mutant FLAG-ELOVL4 decreased at the late stage (Fig. 6). Regarding VGluT2, its expression in the ML showed a trend toward decrease in mice expressing WT ELOVL4 at the late stage. In contrast, its expression in the ML tended to decrease at the early stage and significantly decreased at the late-stage post-onset in mice expressing W246G mutant ELOVL4 (Fig. 6). Since the alteration of VGluT2 expression levels paralleled the severity of motor dysfunction, the reduction of VGluT2 in the ML appears more relevant to the motor dysfunction than that of VGluT1. These findings suggest that the reduction of synaptic inputs from climbing fibers is closely related to motor dysfunction in the SCA34 model mice.

**Figure 6.**
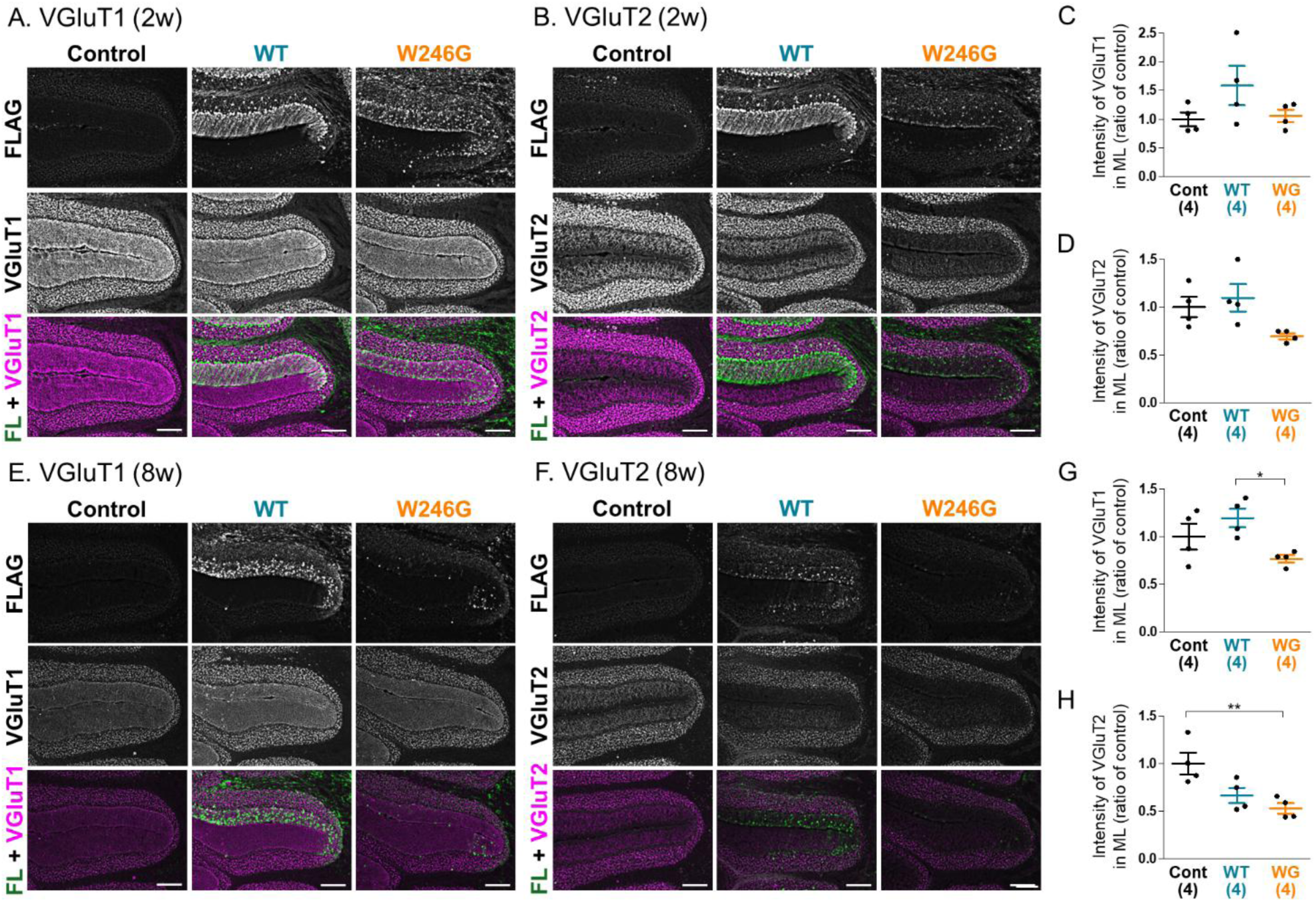
Histological changes in excitatory synaptic inputs to the Purkinje cells in mice expressing wild-type and SCA34 mutant FLAG-ELOVL4 in cerebellar neurons. A,E. Representative immunofluorescence images showing FLAG, VGluT1, and merged FLAG + VGluT1 expression in the cerebellar cortex of control, WT, and W246G mice at 2 weeks (A) and 8 weeks (E) after AAV vector injection. B,F. Representative immunofluorescence images showing FLAG, VGluT2, and merged FLAG + VGluT2 expression in the cerebellar cortex of control, WT, and W246G mice at 2 weeks (B) and 8 weeks (F) after AAV vector injection. Scale bars are 100 μm. C,G. Quantitative analysis of VGluT1 intensity in the ML at 2 weeks (C) and 8 weeks (G) post-injection. D,H. Quantitative analysis of VGluT2 intensity in the ML at 2 weeks (D) and 8 weeks (H) post-injection. Numbers in parentheses in the graphs represent the number of mice assessed per group. ** p < 0.01, *** p < 0.001 (One-way ANOVA).

DCN neurons receive GABAergic inhibitory inputs from cerebellar Purkinje cells (Sastry et al., 1997). Vesicular GABA transporter (VGAT) is localized at the presynaptic terminals of GABAergic neurons. In the DCN, VGAT is strongly co-localized with calbindin, a Purkinje cell marker (Supplementary Fig. 1), suggesting that most VGAT in the DCN is located at the axonal terminals of Purkinje cells. We investigated whether the overexpression of ELOVL4 affects the expression level and pattern of VGAT in the DCN at the early-and late-stages post-onset of motor dysfunction. A significant decline in the total VGAT expression was observed in the DCN of the mice expressing W246G mutant ELOVL4 at the late stage (Fig. 7A,B,D,E). In WT FLAG-ELOVL4-expressing mice, VGAT-positive synaptic terminals outlined most of the somata of FLAG-positive DCN neurons at the early stage (yellow arrow in Fig. 7A,D). Similar VGAT-outlined structures were frequently observed in the DCN of the control mice (yellow arrowhead in Fig. 7A,D). We confirmed that VGAT immunoreactivity outlined NeuN-positive DCN neurons in control mice (Supplementary Fig. 2). Therefore, these VGAT-outlined structures represent the axonal terminals of Purkinje cells. These VGAT-outlined structures were disrupted and significantly decreased in the DCN of W246G mutant FLAG-ELOVL4-expressing mice at the early stage (Fig. 7A,C). At the late stage, the expression of both WT and W246G mutant FLAG-ELOVL4 triggered a prominent decline in the VGAT-outlined structures in the DCN (Fig. 7D,F). Since the total expression of VGAT was not significantly affected in SCA34 model mice at the early stage (Fig. 7B), this structural alteration is likely a result from the disturbance of axonal projection from Purkinje cells. To visualize the Purkinje cell axons, we co-injected AAV vectors to express GFP under the Purkinje cell-specific L7 promoter with vectors to express FLAG-ELOVL4. At the early-stage post-onset of motor dysfunction, the GFP-positive axons in the DCN of WT ELOVL4-expressing mice appeared similar to those of control mice. However, a punctate pattern of GFP-positive axons was observed in the DCN of mice expressing W246G mutant ELOVL4, suggesting that axonal fragmentation was induced by the expression of W246G mutant ELOVL4, leading to the disturbance of axonal projection from Purkinje cells to DCN neurons.

**Figure 7.**
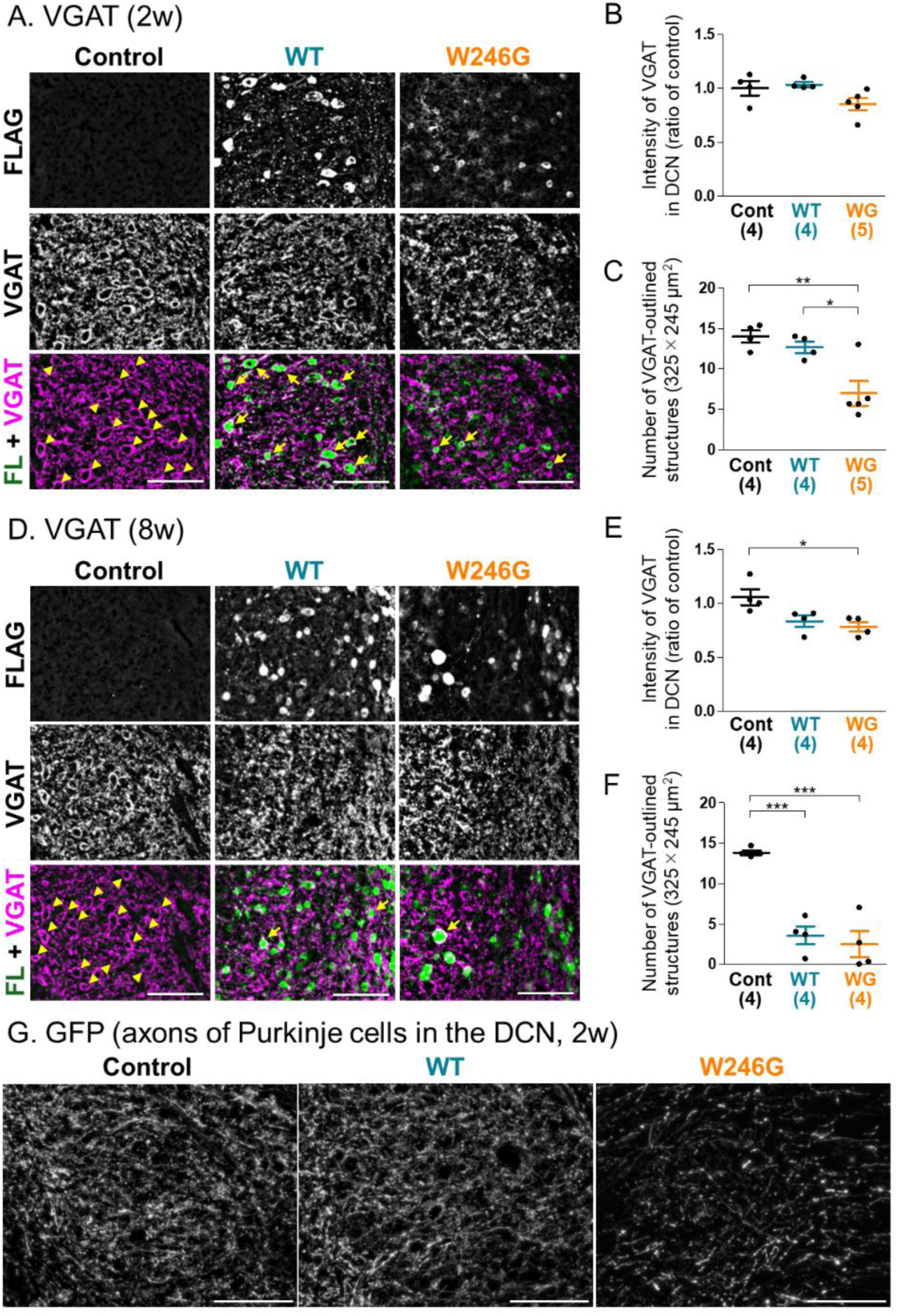
Histological changes in inhibitory synaptic inputs to the DCN neurons in mice expressing wild-type and SCA34 mutant FLAG-ELOVL4 in cerebellar neurons. A,D. Representative immunofluorescence images showing FLAG, VGAT, and merged FLAG + VGAT expression in the DCN of control, WT, and W246G mice at 2 weeks (A) and 8 weeks (D) after AAV vector injection. Yellow arrowheads indicate VGAT-outlined structures; yellow arrows indicate the VGAT-outlined structures and VGAT-outlined neurons expressing FLAG-ELOVL4. Scale bars are 100 μm. B,E. Quantitative analysis of VGAT intensity in the DCN at 2 weeks (B) and 8 weeks (E) post-injection. C,F. Quantitative analysis of the number of VGAT-outlined structures in the DCN at 2 weeks (C) and 8 weeks (F) post-injection. Structures were counted in the same areas shown in A and D. Numbers in parentheses in the graphs represent the number of mice assessed per group. * p < 0.05, ** p < 0.01, *** p < 0.001 (One-way ANOVA). G. Representative fluorescence images of GFP, representing Purkinje cell axons, in the DCN of control, WT, and W246G mice at 2 weeks post-injection. These mice were co-injected with AAV vectors to express GFP in a Purkinje cell-specific manner and either PBS or AAV vectors to express FLAG-ELOVL4. Scale bars are 100 μm.

## Discussion

In the present study, we sought to elucidate the pathogenic mechanism of spinocerebellar ataxia type 34 (SCA34) by establishing a mouse model. We achieved this by directly injecting AAV vectors into the cerebellum to express the missense W246G mutant ELOVL4 specially in cerebellar neurons. Mice expressing mutant FLAG-ELOVL4 in cerebellar neurons developed progressive motor dysfunction starting 2 weeks post-injection compared with control mice injected with PBS (Fig. 1B,C). Since this progressive motor dysfunction resembles that observed in other established SCA model mice (Burright et al., 1995; Seki et al., 2018; Shakkottai et al., 2011; Unno et al., 2012), we successfully in established a functional SCA34 model. Meanwhile, mice expressing WT FLAG-ELOVL4 also displayed mild motor dysfunction, albeit starting later at 4 weeks after AAV vector injection (Fig. 1B,C). A previous immunostaining study on dermal fibroblasts from SCA34 patients with a different missense mutation (L168F), who also presented with skin symptoms, indicated that mutant ELOVL4 forms aggregates within fibroblasts (Beaudin et al., 2020). This suggests that missense mutations in ELOVL4 alter their three-dimensional structures, leading to destabilization and potential contribution to SCA34 pathogenesis. In our previous research on SCA14, while WT PKCγ did not aggregate in cells, PKCγ fused with GFP (PKCγ-GFP) showed mild aggregation and cytotoxicity (Seki et al., 2005). Introduction of various SCA14-related missense mutations into this PKCγ-GFP construct exacerbated the aggregation-prone property, with cytotoxicity correlating directly with the degree of aggregation (Seki et al., 2005). Similarly, the WT ELOVL4 expressed in the current study was labeled with an N-terminal FLAG tag to facilitate detection of the exogenous protein. It is plausible that the FLAG tag might interfere with the three-dimensional structure of WT ELOVL4, causing a milder destabilization than the missense mutation. This could account for the mild SCA-like symptoms observed even with overexpression of wild-type protein. Furthermore, a prior study establishing SCA21 model mice via AAV vector-mediated gene transfer to cerebellar neurons (Seki et al., 2018) also reported motor dysfunction with expression of a FLAG-fused wild-type protein, although it was less severe than that caused by the mutant protein. Collectively, these findings suggest that the FLAG tag itself may possess a property that destabilizes protein structure.

We analyzed neurodegeneration by preparing coronal sections of SCA34 model mice and performing immunostaining for PKCγ (a Purkinje cell marker) and MAP2 (a neuronal marker). Reductions in PKCγ-positive Purkinje cells and ML shrinkage in the cerebellar cortices were both strongly correlated with the motor dysfunction in mice expressing either WT or W246G mutant FLAG-ELOVL4 (Fig. 2). Despite the prominent loss of PKCγ-positive Purkinje cells (to less than 20 % of control mice), the extent of ML shrinkage was relatively small (to 50-90% of control mice). This prominent decrease in PKCγ immunoreactivity relative to the minor ML atrophy initially raised the possibility that the W246G mutant ELOVL4 primarily causes a reduction in PKCγ protein within Purkinje cells rather than outright degeneration. This possibility is supported by reports that PKCγ knockout mice exhibit motor dysfunction (Chen et al., 1995), that PKCγ expression is reduced in cerebellar Purkinje cells of postmortem brains from SCA14 patients with PKCγ missense mutations (Chen et al., 2003), and that PKCγ expression or localization contributes to motor dysfunction in SCA1 model mice (Chopra et al., 2018; Skinner et al., 2001). However, immunostaining for calbindin, another Purkinje cell marker, also showed a drastic decrease in calbindin-positive Purkinje cells (Supplementary Fig. 3). This result strongly suggests that the expression of W246G mutant ELOVL4 eventually triggers Purkinje cell degeneration, leading to progressive motor impairment. The relatively minor effect on ML thickness may be attributable to the persistence of Bergmann glia (Fig. 4), radial astrocytes localized within the ML of the cerebellar cortices (Brockhaus and Deitmer, 2002).

The DCN, which is critical for motor function regulation (Mao et al., 2022; Sastry et al., 1997), integrates inhibitory input from Purkinje cells and excitatory input from climbing and mossy fibers. Neurodegeneration in the DCN, alongside neuronal loss in the cerebellar cortex, has been reported in postmortem brains of patients with SCA2, 3, and 6 (Gierga et al., 2009; Scherzed et al., 2012). In the present study, mutant FLAG-ELOVL4 was expressed in MAP2-positive DCN neurons. This expression triggered a reduction in MAP2-positive neurons and atrophy of cell bodies, effects that correlated with motor dysfunction (Fig. 3). However, in our SCA34 model mice, the degeneration of Purkinje cells was more pronounced than that of DCN neurons at the early-stage post-onset of motor dysfunction, suggesting that Purkinje cells are more vulnerable to mutant ELOVL4. In naturally occurring *Long Evans Shaker* rats, which exhibit motor dysfunction, the degeneration of Purkinje cell axons and the resulting reduced inhibitory input to DCN lead to increased excitability of DCN neurons (Barron et al., 2018). In our SCA34 model mice, the preceding degeneration of Purkinje cells might induce overexcitation and subsequent degeneration of DCN neurons, potentially in addition to the direct toxic effects of the mutant ELOVL4 expressed in DCN neurons themselves.

Histological analysis confirmed that cerebellum-specific expression of mutant ELOVL4 causes the degeneration of both Purkinje cells and DCN neurons. ELOVL4 is an elongase involved in the synthesis of very long-chain saturated fatty acids (VLC-SFAs) and very long-chain polyunsaturated fatty acids (VLC-PUFAs), particularly specializing in VLC-SFA synthesis in the brain (Brush et al., 2010). As a member of the ELOVL family of long-chain fatty acid elongases, ELOVL4 forms homodimers and heterooligomerizes with other family members, and these interactions influence overall fatty acid synthesis (Grayson and Molday, 2005; Okuda et al., 2010). In addition to SCA34, mutations in *ELOVL4* also cause Stargardt macular dystrophy type 3 (STGD3), an autosomal dominant disorder (Logan and Anderson, 2014). STGD3 mutant ELOVL4 loses its C-terminal ER retention motif, which causes it to form dimers with wild-type ELOVL4. This results in the mislocalization of ELOVL4 within cells and negatively affects the enzymatic function of wild-type protein (Grayson and Molday, 2005; Logan and Anderson, 2014). Hippocampal neurons derived from STGD3 model knock-in mice demonstrate increased synaptic release, which can be reversed by the addition of VLC-SFAs (Hopiavuori et al., 2018). It is hypothesized that VLC-SFAs, incorporated into sphingolipids of synaptic vesicle membranes, act as membrane stabilizers, thereby increasing the energy required for the vesicle membrane fusion and suppressing spontaneous synaptic release that is unrelated to action potential firing (Hopiavuori et al., 2018). Further supporting the role of lipid dysfunction, lipid analysis of the retina and skin of W246G homozygous knock-in rats ‒‒carrying the same mutation used in this study— showed a selective impairment of VLC-SFA synthesis (Agbaga et al., 2020). More recently, cells expressing two types of SCA34 missense mutant ELOVL4 (L168F, W246G) were shown to produce altered types of VLC-PUFAs and a significantly reduced amount of VLC-SFAs compared to cells expressing wild-type ELOVL4 (Gyening et al., 2023). As previously discussed, missense mutations in ELOVL4 may cause destabilization of its three-dimensional structure. Consequently, the mutant ELOVL4 could potentially disrupt the structure, enzymatic activity, and intracellular localization of the wild-type protein, leading to an alteration in VLC-SFA synthesis capacity and VLC-PUFA synthesis characteristics. In the SCA34 model mice used here, expression of W246G mutant FLAG-ELOVL4 in cerebellar neurons might impair VLC-SFA synthesis. This impairment could cause abnormal neurotransmission due to excessive synaptic release, eventually leading to atrophy and loss of Purkinje cells and DCN neurons through excitotoxicity or excessive inhibition of neuronal excitation. Additionally, since VLC-PUFAs are known to affect cell membrane structure and fluidity (Cheng et al., 2022), the changes in VLC-PUFA types resulting from mutant ELOVL4 expression may have a more significant impact on Purkinje cells, which possess highly developed dendrites and larger cell membrane surface areas. This heightened sensitivity could explain the greater vulnerability of Purkinje cells compared to DCN neurons and interneurons.

Investigation of glial cell activation revealed that microglial activation (indicated by increased Iba1 expression) occurred in both the cerebellar cortices and the DCN in SCA34 model mice prior to the onset of motor dysfunction (Fig. 4,5). Microglial activation is known to be detrimental to neurons through mechanisms including phagocytosis of Purkinje cell dendrites, glutamate-mediated excitotoxicity, inflammatory cytokine release, and oxidative stress (Childs et al., 2021; Kavetsky et al., 2019). Thus, in our SCA34 model, microglial activation in the cerebellar cortex likely contributes to Purkinje cell degeneration and plays a crucial role in the onset of motor dysfunction. Astrocyte activation (indicated by increased GFAP expression), conversely, was not detected prior to motor dysfunction. It only became apparent in the cerebellar cortex and the DCN after onset and progressed in severity parallel to the worsening motor symptoms (Fig. 4,5). Early glial cell activation, either before or concurrent with the onset of motor dysfunction, has been similarly observed in other SCA models (Aikawa et al., 2015; Cvetanovic et al., 2015; Ohta et al., 2021; Seki et al., 2018). Moreover, optogenetic and chronic activation of cerebellar Bergmann glia in mice has been shown to induce Purkinje cell loss and ML shrinkage (Shuvaev et al., 2021). Therefore, astrocyte activation is likely a contributing factor to cerebellar neurodegeneration, including the observed Purkinje cells loss.

Our findings suggest a sequence where preceding microglial activation triggers and subsequently enhances astrocyte activation. However, the mechanism by which SCA34 mutant ELOVL4 expression in neurons induces microglial or astrocyte activation remains unclear. We have previously demonstrated that a decline in chaperone-mediated autophagy (CMA), a component of the autophagy-lysosome protein degradation system, is linked to the pathogenesis of several SCAs (Sato et al., 2021; Seki et al., 2018, 2012). D-cysteine, which is converted to hydrogen sulfide in the cerebellum (Shibuya et al., 2013), has been shown to alleviate dendritic shrinkage caused by various SCA-related proteins and prevent the onset of motor dysfunction in SCA model mice by activating CMA in Purkinje cells (Ohta et al., 2021; Ueda et al., 2022). Furthermore, we have demonstrated that reduced CMA activity enhances the release of exosomes, a form of secretory vesicles (Oshima et al., 2019). In models of traumatic brain injury (TBI), neuron-derived exosomes containing miR-21-5p are phagocytosed by microglia, inducing an inflammatory phenotype, cytokine secretion, and subsequent neuronal apoptosis (Yin et al., 2020). Taken together, these findings suggest that SCA34 mutant ELOVL4 may impair CMA in cerebellar neurons, thereby promoting exosome secretion that leads to microglial activation via neuron-derived exosomes. Our prior work showing that a selective decline of CMA activity in the cerebellar neurons triggers both microglial and astrocyte activation before motor dysfunction (Sato et al., 2021) further supports this hypothesized link between decreased neuronal CMA activity and early glial activation. miR-21-5p is known to suppress the expression of CMA-related proteins LAMP2A and Hsc70 (Alvarez-Erviti et al., 2013). Its level is also increased in patients with Parkinson’s disease and Friedreich ataxia (a recessive ataxic disorder) (Dantham et al., 2018). However, changes in miR-21-5p expression have not yet been elucidated in SCA patients or animal models. Further studies are necessary to clarify the detailed mechanisms of this early glial activation. Immunohistological analysis of the cerebellar cortex revealed that the expression of VGluT1, a marker for the presynapses of parallel fibers (Hioki et al., 2003; Mao et al., 2022), was unaffected in SCA34 model mice at the early-stage post-onset but significantly declined in at the late stage (Fig. 6). Since degeneration of Purkinje cells was already evident at the early stage, this reduction of VGluT1 is likely a consequence of the overall decrease in Purkinje cells. In contrast, VGluT2, which is expressed in the presynapses of climbing fibers projecting to the cerebellar cortex, showed an early tendency to decrease in the cerebellar cortices of SCA34 model mice (Fig. 6). These results suggest that the synaptic transmission from climbing fibers to Purkinje cells is more vulnerable to the toxicity of SCA34 mutant ELOVL4 expressed in cerebellar neurons. A similar observation is seen in SCA1 model mice, where synaptic activity between climbing fibers and Purkinje cells declines at the early stage of motor dysfunction, while parallel fiber activity remains stable until the further disease progression (Barnes et al., 2011). Furthermore, the nicotinamide antagonist 3-acetylpyridine causes ataxia in rats by inducing degeneration of inferior olive neurons, which are source of climbing fibers (Llinás et al., 1975). Therefore, loss of synaptic inputs from climbing fibers may be an early contributor to motor dysfunction in SCA34.

In the DCN, VGAT is localized within Purkinje cell axon terminals, where it surrounds the somata of DCN neurons (Carulli et al., 2020) (Supplementary Fig. 2). At the early-stage post-onset, while the overall VGAT expression level remained unchanged, the localization of VGAT around the DCN neuron somata was significantly disturbed in SCA34 model mice (Fig. 7). We proposed two possible explanations for this VGAT mislocalization. One is the axonal damage in Purkinje cells. The expression of mutant ELOVL4 in Purkinje cells may trigger axonal damage. Indeed, we observed axonal fragmentation at the early-stage post-onset in SCA34 model mice (Fig. 7G). Previous research showing that Purkinje cell-specific knockdown of Atg7 (essential for macroautophagy) leads to axonal dystrophy and motor dysfunction (Komatsu et al., 2007) highlights the importance of the autophagy-lysosome system in maintaining Purkinje cell axons. As we propose that CMA impairment is a common pathogenic pathways in SCAs (Sato et al., 2021; Seki et al., 2018, 2012), CMA impairment in our SCA34 model might similarly disrupt the homeostasis of axonal proteins and trigger axonal damage. However, the AAV vector injection used here only transduces a fraction of Purkinje cells projecting to the DCN (Ohta et al., 2021). Despite many VGAT-positive terminals originating from untransduced Purkinje cells, the change in VGAT distribution was observed around most DCN neurons in the examined area (Fig. 7). The second possible explanation for this VGAT mislocalization is the alteration of the DCN neurons via microglial activation. The widespread mislocalization suggests that an alteration in the DCN neurons themselves may be responsible, potentially triggered by the observed microglial activation. Perineuronal nets (PNNs), composed of extracellular matrix molecules (chondroitin sulfate proteoglycans, hyaluronan, etc) (Carulli et al., 2006), surround DCN neuron somata (Carulli et al., 2020) and regulate synapse stability and neural plasticity (Reichelt et al., 2019). Critically, a recent report demonstrated that microglial activation triggers PNN loss (Liu et al., 2024).

Since VGAT-positive terminals are colocalized with PNNs in the DCN (Carulli et al., 2020), the early microglial activation detected before the onset might cause PNN loss and subsequently disturb the precise outline structure of VGAT around the DCN neuron somata in the SCA34 model mice. Regardless of the underlying cellular mechanism, the disturbed VGAT localization indicates that a loss of normal GABAergic inputs from Purkinje cells to DCN neurons, which likely contributes to the motor dysfunction and subsequent DCN neuron degeneration observed in the SCA34 model mice.

Previous studies using a W246G mutant ELOVL4 knock-in rat model of SCA34 demonstrated impaired synaptic plasticity (parallel/climbing fibers to Purkinje cells) and reduced dendritic spine density of Purkinje cells, leading to progressive motor impairment (Nagaraja et al., 2023, 2021). However, the rat model did not exhibit cerebellar neurodegeneration (Fessler et al., 2024). In contrast, clinical reports of SCA34 patients confirm cerebellar neurodegeneration and cortical shrinkage (Beaudin et al., 2020; Ellezam et al., 2023; Ozaki et al., 2021b, 2019). In the present study, we successfully established SCA34 mouse model that exhibits progressive motor dysfunction accompanied by the degeneration of both Purkinje cells and DCN neurons. Furthermore, we have revealed several possible pathogenic mechanisms of SCA34, including (1) early activation of microglia, (2) differential synaptic vulnerability (climbing fiber input to Purkinje cells), and (3) disturbance of the synaptic structure between Purkinje cells and DCN neurons (VGAT mislocalization). Given that this novel SCA34 model shares key features observed in human patients and other SCA models (Aikawa et al., 2015; Cvetanovic et al., 2015; Ohta et al., 2021; Seki et al., 2018), and exhibits a rapid progressive motor dysfunction compared to some other SCA models (Ohta et al., 2021; Seki et al., 2018), it represents a valuable tool for screening therapeutic and preventive drugs for SCA34 and for various other SCAs.

## CRediT authorship contribution statement

**Yuri Morikawa-Yujiri**: Investigation, Visualization, Data curation, Formal analysis, Writing – original draft, **Kensuke Motomura**: Investigation, Visualization, **Ayumu Konno**: Resources, Funding acquisition, **Natsuko Hitora-Imamura**: Validation, Supervision, **Yuki Kurauchi**: Methodology, Validation, Supervision, **Satohiro Masuda**: Writing – review & editing**, Hirokazu Hirai**: Resources, Funding acquisition, **Hiroshi Katsuki**: Resources, Methodology, Validation, Supervision, Conceptualization, Writing – review & editing**, Takahiro Seki**: Conceptualization, Data curation, Formal analysis, Visualization, Supervision, Funding acquisition, Validation, Methodology, Project administration, Writing – review & editing

## Data availability statement

Data generated or analyzed in this study are available on reasonable request by contacting the corresponding author.

## AI declaration statement

The authors utilized Gemini (Google) to improve the English language quality and clarity of the manuscript. The authors alone are responsible for the content and scientific integrity of the final submission.

