## Supplementary figure 1, Supplementary figure 2, Supplementary figure 3 for "Establishment of spinocerebellar ataxia type 34 model mice accompanied by early glial activation and degeneration of cerebellar neurons"

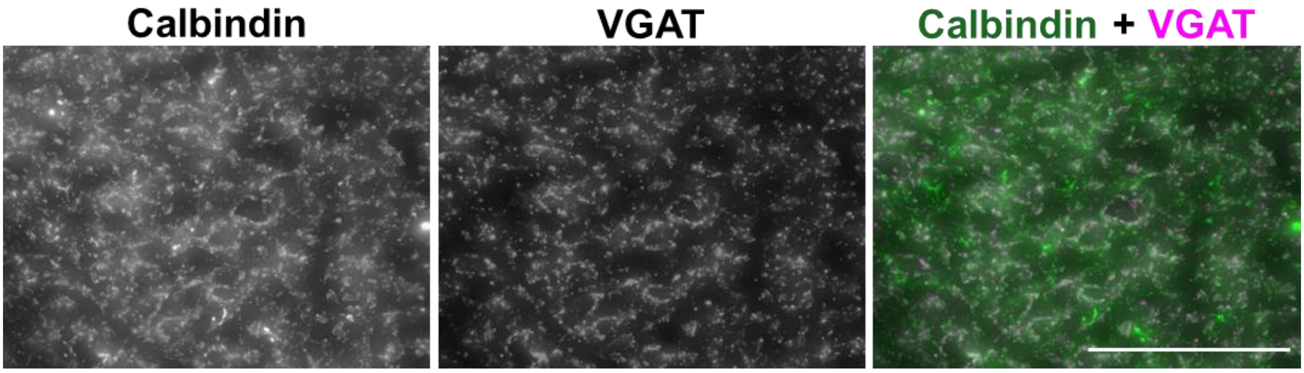


**Supplementary figure 1. Localization of VGAT to the axonal terminals of Purkinje cells.**

Representative immunofluorescence images showing calbindin (a Purkinje cell marker), VGAT, and merged calbindin + VGAT expression in the DCN of the control mouse. Scale bar is 100 μm.

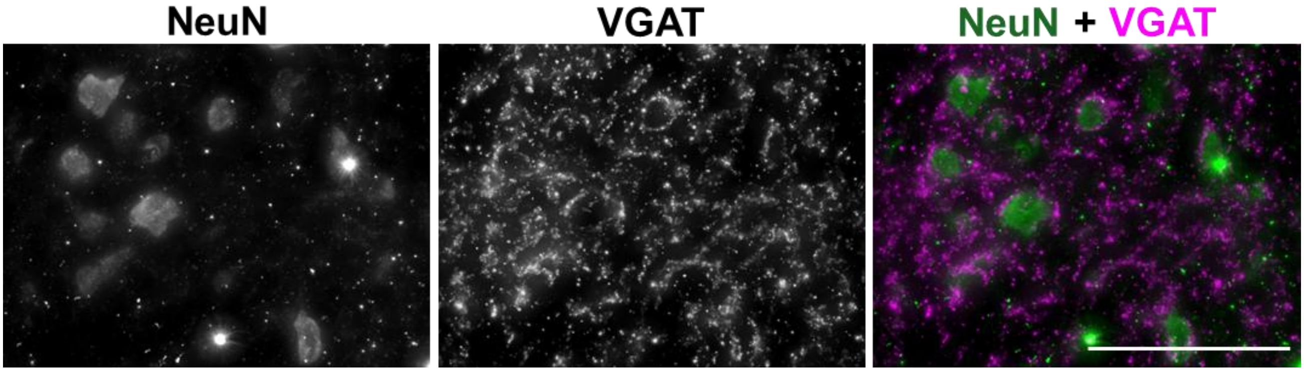


**Supplementary figure 2. Localization of VGAT around the DCN neurons.**

Representative immunofluorescence images showing NeuN (a neuronal marker), VGAT, and merged NeuN + VGAT expression in the DCN of the control mouse. Scale bar is 100 μm.

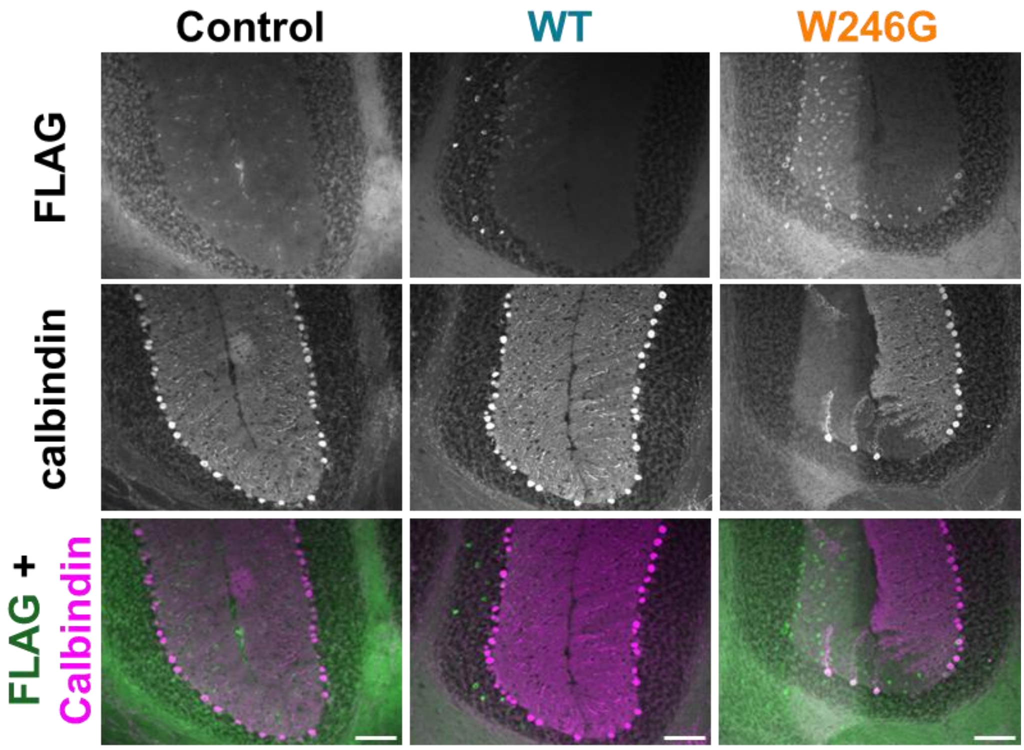


**Supplementary figure 3. Survival of Purkinje cells in the cerebellar cortices of mice expressing wild-type and SCA34 mutant FLAG-ELOVL4.**

Representative immunofluorescence images showing FLAG, calbindin, and merged FLAG + calbindin expression in the cerebellar cortices of the control, WT, and W246G mice at 2 weeks after AAV vector injection (at the early-stage post-onset). Scale bars are 100 µm.
